# Prokaryotic SPHINX sequences are conserved in mammalian brain and participate in neurodegeneration

**DOI:** 10.1101/593954

**Authors:** Klara Szigeti-Buck, Laura Manuelidis

## Abstract

A new class of circular mammalian “SPHINX” DNAs, represented by the 1.8 and 2.4kb elements, were discovered in highly infectious cytoplasmic particles isolated from Creutzfeldt-Jakob Disease (CJD) and scrapie samples. These DNAs contained replication initiation sequences (REPs) with ∼70% homology to Acinetobacter phage REP segments. Antibodies against REP peptides from the 1.8kb DNA highlighted a 41kDa protein (spx) on Western blots, and in-situ studies revealed its tissue and cell-type specific expression, e.g., in pancreatic islet cells, keratinocytes and kidney tubules but not pancreatic exocrine cells, alveoli, and striated muscle. An intense spx signal in oocytes implicated maternal inheritance of the SPHINX 1.8 genome, a feature known only for bacterially derived mitochondrial DNA in mammals. To determine if spx concentrated at specific neurons and synapses, and also maintained a conserved pattern of architectural organization in mammals, we evaluated mouse, rat, hamster, Guinea Pig and human brains. Most outstanding was the cross-species concentration of spx in huge excitatory synapses of mossy fibers and small internal granule neuron synapses, the only excitatory neuron within the cerebellum. This synaptic localization was also demonstrable at the ultrastructural level. Vacuolar loss of these synaptic complexes, thinning of the internal granule cell layer, and pathological fibrillar spx accumulations within Purkinje neurons were obvious in CJD Guinea pig brains. In rats, these fibrillar changes marked hippocampal Pyramidal neurons and preceded prion protein misfolding. Spx may define different causes or processes of neurodegeneration. The evolutionary origin, persistence and modulation of SPHINX 1.8 opens an unexpected chapter in mammalian symbiosis.

## INTRODUCTION

The causes of late-onset neurodegeneration can be complex and heterogeneous. Unrecognized environmental, toxic, and microbial agents may initiate and contribute to the development of major common diseases such as Alzheimer’s Disease (AD). The current strong focus on end-stage amyloid and a few other fibril-forming proteins has diverted investigation of unsuspected microbial causes of late-onset neurodegeneration (1, 2) and these have some precedent. Influenza viruses have caused late onset neurofibrillary tangles (NFTs) in Parkinson’s neurodegeneration in a “hit and run” pattern, as in the 1918 epidemic, and activated latent myxo-paramyxoviral infections can induce NFTs (3). NFTs are also prominent in AD and in post-traumatic dementia, and this NFT phenotype fails to discriminate viral, traumatic, and other initiating causes of AD. Targeting environmental infectious pathogens can have major beneficial health effects for neurodegenerative diseases. Indeed, epidemic “mad cow” disease, a transmissible encephalopathy (TSE), was dramatically reduced simply by removing infectious animals and their products from the food chain. We have worked extensively with TSEs because these agents cause stealth infections that can remain quiescent in lymphoreticular cells for many years without inciting neurodegenerative sequelae. The essential molecular components of these infectious agents remain undefined (4, 5), and they present an intriguing evolutionary mystery. As other latent viruses, TSE agents do not provoke an acute lymphocytic reaction but they can induce early innate immune responses as they begin to exponentially replicate in brain (1). These host responses include dendritic cell and microglial changes with neuronal vacuolization that occurs well before any pathological prion protein (PrP) appears in brain (6-8). Microglia are also prominent players in other late-onset neurodegenerations. Elucidating intrinsic TSE agent molecules can help prevent TSEs and offer new insights in other neurodegenerative diseases of unknown cause.

It is widely believed that host PrP, without nucleic acid, spontaneously misfolds to become the causal infectious TSE agent (9, 10). Prion protein misfolding has also been claimed to encode all the different TSE agent strains (11). These infectious strains produce vastly different incubation times, species transmissibility and regional pathology, yet they elicit identical PrP misfolded band patterns in brain (12). PrP bands change when TSE agents are propagated in different cell types, but the strain does not. Even after a year of propagation in monotypic culture, cell type specific changes in PrP misfolding do not alter the strain properties (13). Some TSE agents can also interfere with infection by a second TSE strain whereas others can add their infectivity, and this viral behavior is not predicted by any specific PrP misfolding (14-16). Two reports claim recombinant PrP (recPrP) itself, without mammalian nucleic acids, can become infectious when misfolded into fibrils in vitro (17, 18), but thousands of experiments elsewhere have failed to reproduce this result (19-21). Moreover, highly infectious brain particles of viral size (∼20nm diameter), comparable to those seen only in infected brain and cell sections (22), have lacked detectable PrP by deep proteomic sequencing (4). In contrast, highly infectious cytoplasmic particles always contain nucleic acids of >500nt, and particle infectivity and nucleic acids are both destroyed together by nucleases while PrP remains unaffected (5). The discovery of new class of circular SPHINX DNAs derived from a logical interrogation for a protected TSE genome.

Because a ∼20nm infectious particle can accommodate a circular DNA genome of 1-5kb (23), nucleic acids protected in cytoplasmic particle fractions were analyzed using rolling circle amplification (23). At least four related circular DNA elements were appreciable in brain and cultures. Two of these circular elements with REP motifs (of 1.758 and 2.36kb) were completely sequenced, along with copurifying circular mitochondrial DNA of 16kb (23). Whereas mitochondrial sequences had a 100% homology with those in the database, both REPs had only an ∼70% significant homology (e^-105^) to small REP segments of an Acinetobacter phage virus, e.g., 676/976 bases. Extensive tests failed to show these new circular DNAs were in any chemicals or reagents used. Although both of SPHINX DNAs were highly enriched in infected preparations, they subsequently were identified in noninfectious material by both PCR and antibody studies, and their origin and representation were unclear, as was their potential involvement in neurodegeneration. The 1.8kb DNA was preferentially targeted for further study because it contained a classical iteron initiation repeat. It was also independently isolated and verified in Europe in 3 bovine sera clones that were 100% homologous to our 1.758nt sequence. An association with neurodegeneration was also substantiated because two multiple sclerosis brain samples contained 1.766nt sequences with a 98% nucleotide identity to SPHINX 1.8 (24).

We generated REP-specific antibodies to first delineate the expression of the 1.8kb sequence in different tissues and cells. Two rabbits both produced antisera against a REP peptide not present in the mammalian data base. Both sera labeled a non-glycosylated 41kDa protein band, close to its predicted 38kDa size, and labeling was completely blocked by the REP peptide. The highest specific titer serum was affinity purified. On Western blots this IgG showed a strong 41kDa signal at 1:5,000 dilution that was completely blocked by the REP peptide in Western Blots, cell culture, and tissue sections as shown previously (2, 25). Because selected neurons, such as large anterior motor horn neuros appeared to receive synaptic boutons with high concentrations of spx by light microscopy, it was important to substantiate this at the ultrastructural level. The cerebellum showed the most striking concentration of synaptic spx in a highly organized and selective pattern as reported here. Dendritic claws of excitatory internal granule cell (GC) neurons (26) receiving large pontine excitatory synapses showed strong spx accumulations, and these GC cells synchronize temporospatial coordination (27) that is critical for survival. Each GC cell axon then ascends through the molecular layer (as climbing parallel fibers) and branches to synapse with as many as 100 Purkinje cell dendrites. As demonstrated here their spx positive axonal boutons surrounded Purkinje cell dendrites in the molecular layer, and this feature was also conserved in different species. Additionally, although Western blots, including those from PrP knockout mice, failed to show any quantitative spx differences in CJD and scrapie infected brains versus controls (25), evaluation of cerebellar tissue in situ studies here demonstrated novel spx neurodegenerative changes in the cerebellum as well as in limited studies of the hippocampus, another region rich in excitatory glutamate synapses. These changes included abnormal collections of spx in the Purkinje perikaryon and obvious loss of internal granule neurons with their dendritic synapses. Tracer studies in AD have also shown unappreciated degeneration of GC neurons unrelated to β-amyloid neuritic plaques (28) and spx analysis can be informative in AD. On an evolutionary scale, spx elements represent the tip of the iceberg of new and unsuspected environmental symbiotes that can cause or propel neurodegeneration.

Finally, from a broader evolutionary perspective, SPHINX DNAs appear to be the only viral phage REP reported to reside in the cytoplasm of mammalian cells. The only other protected prokaryotic genome maintained in the cytoplasm of mammals is the 16kb mitochondrial DNA that was acquired early in evolution by endosymbiosis (29). REP circular DNAs fall within the broad class of circular single stranded “CISS” viruses that are found within many plant and invertebrate cells (30), and it is surprising that comparable prokaryotic elements have not been detected previously in mammals, possibly because of their relatively low cytoplasmic copy number. In contrast human endogenous retroviruses (LINES), first isolated and sequenced here in 1982 (31), derive from full length retroviruses and constitute ∼2% of nuclear chromosomal DNA, a huge amount. Along with other endogenous retroviral sequences (ERVs) these elements were retrotranscribed and inserted as DNA early in evolution. During their long infectious and symbiotic interactions with mammals, LINES and other ERVs have developed unique sequence signatures that allow them to define functionally cohesive megabase gene regions in chromosomes (32, 33), promote tissue-specific patterns of gene expression (34), modulate innate immunity (35) and attack competitor species (33). Although the function of SPHINX 1.8 or other members of this family is unknown, this DNA may modify its host, and also be sculpted or altered by its host over time. The synaptic concentration of SPHINX 1.8 as well as other tissue-specific features give credence to an evolving non-random symbiotic modulation.

The studies here define potential new functions for spx in neuronal subsets, and also demonstrate the participation of spx in neurodegenerative processes that occur long before abnormal prion protein accumulates.

## RESULTS

### Light microscopy of spx

Cerebellar folia have a well defined and conserved architecture in mammals with 3 distinct macroscopic layers. This layered architecture facilitates unambiguous identification of specific neuronal cell types with known physiological functions. Different patterns of spx protein are detected in each layer. Alkaline phosphatase labeling was used to detect a red spx signal by transmitted light microscopy and also showed a corresponding intense red fluorescence by epifluorescence as depicted respectively in Figs. 1A, B and C, D. At low magnification A, a section of human cerebellum, the outer molecular layer (m) shows intense red signal throughout. This layer is separated from the internal granule cell layer (g) by a row of Purkinje neuronal cell bodies (P) as shown in C and D. Each Purkinje cell body receives inputs via its complex dendritic arborizations within the molecular layer, and each Purkinje dendritic tree receives excitatory synapses from thousands of granule cell (GC) axons. The GC cell bodies and their stubby branched dendrites reside in the internal granule cell layer (g) where their small neuronal nuclei are stained dark blue in C. Note the many separate and intense discrete red bodies of 2-6μm in this granule cell layer. These are more dramatically accentuated by fluorescence microscopy in D where the perikaryon and nuclei show minimal spx fluorescence. The large congregation of spx signals (rosettes) in this layer correspond to mossy fiber synapses that decorate the GC dendritic claws (at arrows). Mossy neuron synapses, as those of GC neurons, are excitatory and their perikarya reside far away in pontine nuclei of the spinal cord. In Fig. 1E, a guinea pig cerebellum shows the same large spx positive GC clusters as seen in the human section. It also shows many discrete synapse-sized red boutons around minimally labeled processes cut longitudinally in the molecular layer. A plethora of these synapses are excitatory GC axon synapses from their climbing and branched parallel axon branches. Each GC axon can synapse on as many as 100 Purkinje cell dendrites. This section also cuts through a normal Purkinje cell body (P) that shows a much weaker cytoplasmic spx signal. Fig. 1E also shows a stellate interneuron in the molecular layer with its dendrite surrounded by intense spx signals (arrows). These stellate neurons receive excitatory GC inputs as well as inhibitory inputs from other neurons.

**FIGURE 1:**
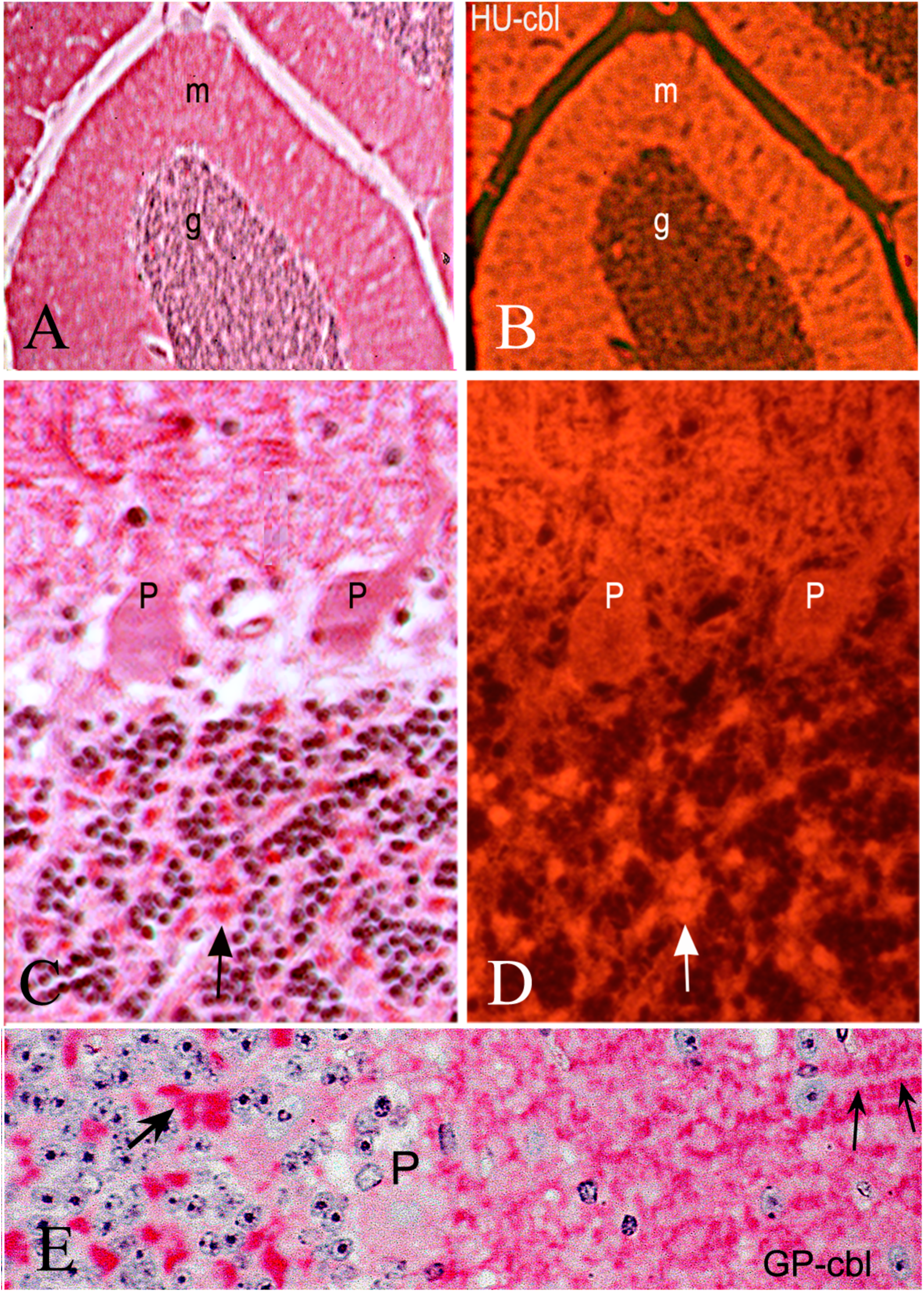
Transmitted light with counterpart fluorescent epi illumination of red spx signals at low (A/B) in low and higher magnification (C/D) of human cerebellum. Outer molecular layer (m) with blue stained nuclei in internal granule cell layer (g). Purkinje cells (P) with dendrites in the outer molecular layer with abundant synapses, many of which come from small granule neuron mossy fiber axons. Arrow in C/D show complex spx+ large excitatory synapses from distant neurons in the pons. E shows Guinea Pig (GP) cerebellum with same spx architectural and synaptic pattern. Large arrow again shows mossy neuron synaptic complex and small arrows point to spx+ boutons on a stellate neuron dendrite in the molecular layer with low spx signal. The Purkinje neuron perikaryon P is also has low spx.

The overall architectural pattern of spx, with high concentrations in rosette synapses on GC dendrites and in numerous GC synapses surrounding dendrites in the molecular layer was consistently seen in all mammals examined (human, guinea pig, hamster, rat and mouse). As in previous studies in peripheral tissues using procedures to enhance tissue penetration (2), neither 2M GdnHCl that can loosen protein binding to nucleic acids, nor other unmasking procedures, altered the pattern and specificity of spx labeling, and this was true even in ultrastructural studies here (see below). The strongly positive mossy fiber synapses in the GC layer contrasts with minimally labeled cytoplasm of GC cell bodies and other adjacent fibers (see C & E). The weak labeling of the GC perikaryon also contrasts with their synapses in distant regions of the molecular layer. Similarly, the strong concentration of spx in mossy fiber synapses, but not their axons traveling through the cerebellar white, matter implicates inapparent transport of spx to synapses and/or synthesis of spx within these synaptic regions. It remains to be determined if SPHINX 1.8 DNA circles are clustered, transcribed and translated within synaptic regions. In any case, synaptic spx is in a perfect position to enforce both rapid and long term plasticity that is critical for the many complex integrative functions of the cerebellum.

To better resolve the structure of spx positive clusters we first used confocal microscopy deconvolution and reconstruction on ∼6μm sections that allow excellent penetration of the vast majority of cells and their organelles. This analysis delineated the discrete spx positive boutons on Purkinje cell dendrites within the molecular layer (m). Fig. 2 shows a sample of a human cerebellum section in two 3D rotations. One large diameter Purkinje cell apical dendrite (between small arrows) is surrounded by discrete round spx bodies consistent with the small axonal synapses from many GC neurons. There is also strong spx labeling of small round boutons surrounding the Purkinje cell body (large arrow) whereas spx labeling in the cytoplasm of the Purkinje perikaryon is relatively weak. The granule cell layer (g) contains much larger spx synaptic signals, and again, surrounding neuronal and glial processes as well as GC cell bodies display minimal signal.

**FIGURE 2:**
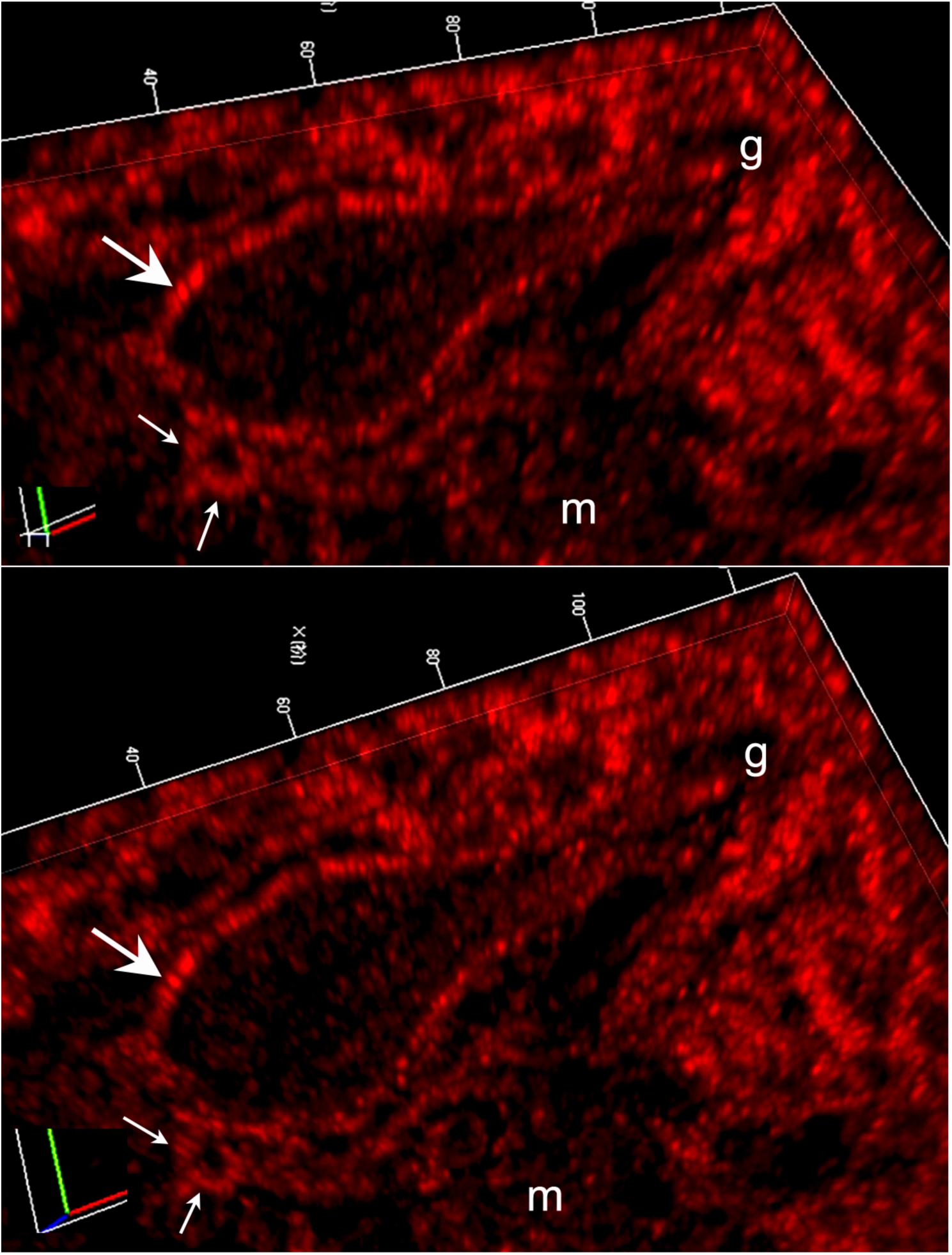
**A** deconvolved fluorescent stack in two close 3-D rotation views of human cerebellum show cross-section of a large Purkinje perikaryon, with low spx signal studded with spx+ round boutons consistent with synapses as at large arrow. Its dendrite is also surrounded by spx+ boutons in the molecular layer and prominent larger spx+ synapses in the internal granule layer (g).

### Ultrastructural concentrations of spx in specific synapses

Even better resolution was obtained in ultrastructural studies. The specific labeling of mossy fiber excitatory synapses (also known as rosettes) around GC dendritic claws was confirmed unequivocally by immuno-electron microscopy. As shown in Fig. 3 of mouse cerebellum, peroxidase precipitates delineate spx antibodies that are prominent in the huge mossy fiber rosettes around the GC claws. In A, two large positive processes (large arrows) are strongly labeled with a smaller positive process between them (smaller arrow near vessel V) in this preparations pretreated with 2M GdnHCl for enhanced permeabilization. These 3 positive processes are consistent with a mossy fiber rosette where only a few small diameter irregular connecting processes are apparent (small arrows) in thin section. Notably the leftmost process is 4μm in length, and the span of this synapse is 8.6μm wide if the large rightmost process is included. This size corresponds to the large synapses seen by light microscopy above, as well as by other classical studies demonstrating a fundamentally conserved fine structure and organization of mossy fiber synapses in species from amphibian through mammals (36). Note the many mitochondria stuffed into these broad terminals, another characteristic of mossy neuron synapses. In Fig. 3B, where freeze-thawing was used for permeabilization, the spx positive processes again demonstrate all the characteristics of mossy fiber inputs (36); multiple synaptic vesicles are seen in the cytoplasm, this process makes many synapses on unlabeled small granule cell dendrites, as at arrows, and the granule cell dendrites also display post synaptic densities at these synapses (arrows). Again, mitochondria are even more abundant in this terminal, a feature that may relate to a replicative and synthetic ability of mitochondria (and perhaps SPHINX 1.8 DNA) at sites distant from their perikarya in chicks and mammals (37, 38).

**FIGURE 3:**
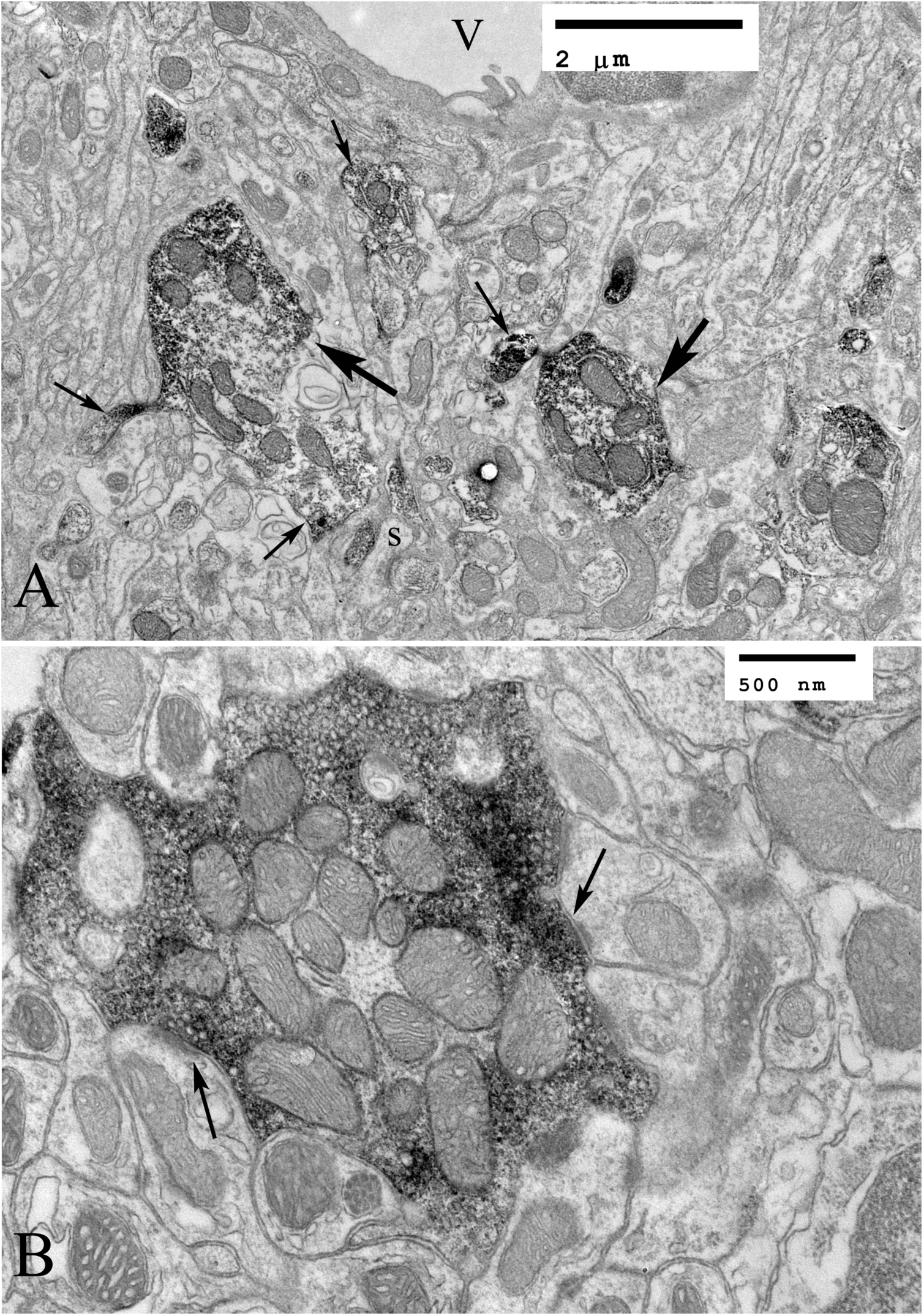
Mouse cerebellum. Ultrastructural localization of spx in mossy neuron excitatory synapses (dense precipitates in complex large processes at arrows in A. V is. vessel. Permeabilized with 2M GdnHCl. B shows another labeled process in higher power with characteristic numerous small vesicles that is also packed with mitochondria. Arrows point to post synaptic densities of the delicate granule cell dendrites innervated by this mossy fiber’s synapses. This preparation was permeabilized only by freeze-thawing (see Methods)

### Spx in other excitatory synapses and cell bodies

The high concentration of spx at identified excitatory mossy neuron glutamate synapses does not exclude its localization at other non-excitatory synapses in the cerebellum and elsewhere. Nor does it exclude spx in the cell body of selected other neurons. Although characterizing all spx positive synapses in the cerebellum with double labeling for functional discrimination is not in the realm of this paper, we evaluated spx in CA3 of hippocampus ultrastructurally for more information. This region also has distinct anatomical layers that contain other types of mossy fiber neurons and small dentate neurons. It is responsible for complex integrative physiology in memory and can also be a target of AD early neurodegeneration. CA3 again revealed labeling of specific synapses, many of which are known to have excitatory properties by light microscopy (data not shown) and this was confirmed ultrastructurally. Fig. 4A shows a section through the CA3 with four spx positive post-synaptic termini that are consistent with those containing excitatory glutamate receptors. Three of these synapses have an ovoid shape and are ∼0.4μm in diameter, characteristics of recurrent synapses that transmit excitatory glutamate from CA3 pyramidal cells to other pyramidal neurons and to interneurons (39). These synapses, as glutamate excitatory cerebellar synapses, have a key role in the synchronous physiology of the hippocampus and integrative memory. For reference, this section also contains three myelinated axons, e.g., at A, that show no spx signal, and this corresponds to the lack of spx in myelinated axons elsewhere. Fig. 4B from the mouse cerebral cortex permeabilized by freeze thawing also shows selective spx labeling of small ovoid synapses, comparable to those in CA3. We also examined large ganglia of the sympathetic-parasympathetic system in the peritoneum, a completely different monotypic group of large neurons. These large neurons, unlike cerebellar Purkinje neurons and hippocampal Pyramidal neurons, clearly display intense spx signal around their nuclei (Fig 5, arrows), within tubal structures likely to be endoplasmic reticulum, and at the membrane. This demonstrates that the intracellular localization of spx is highly specific for different neuronal cell types where it may subserve different functions. In peripheral tissues the intracellular localization of spx is also cell-type specific (2).

**FIGURE 4:**
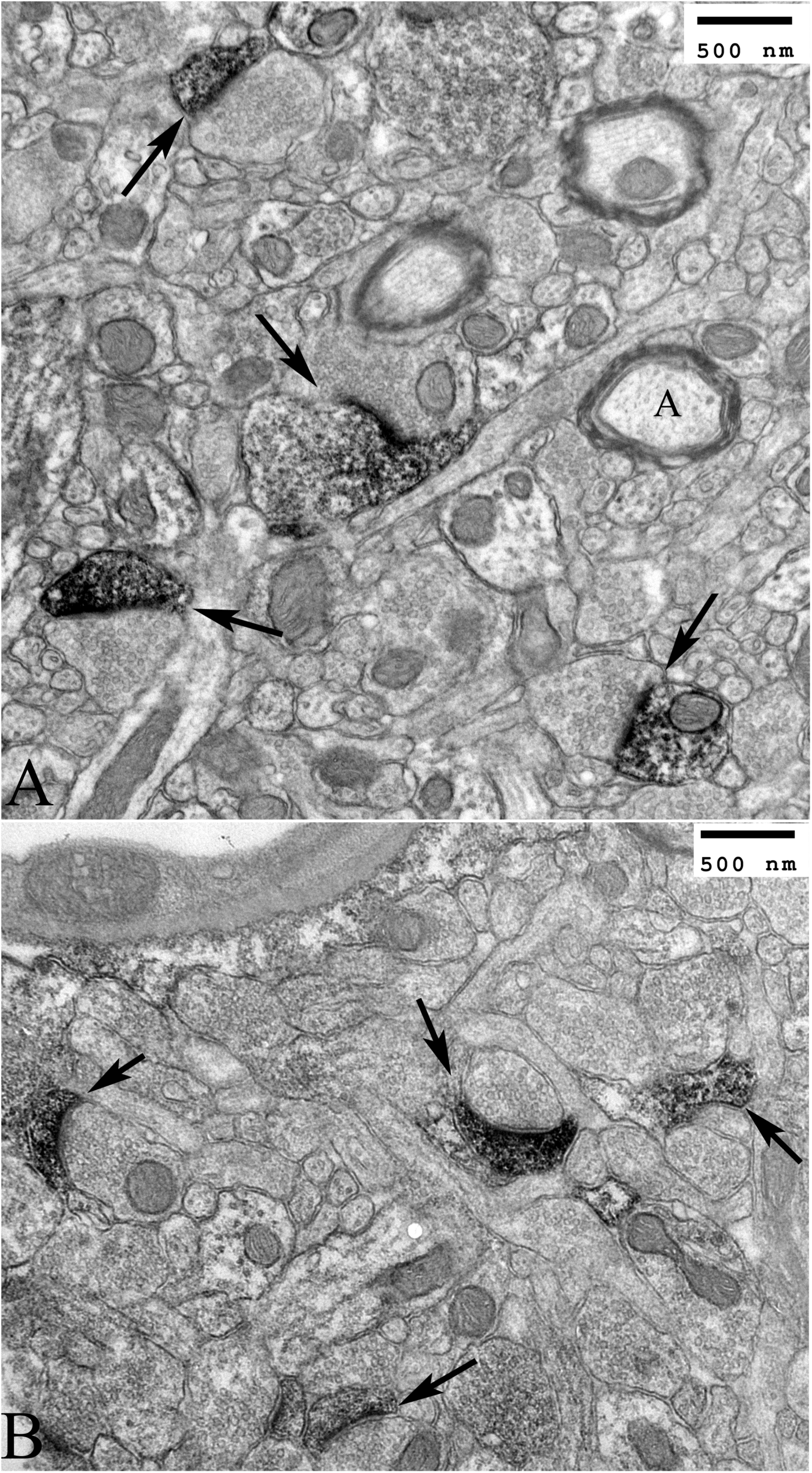
**A:** Mouse hippocampus CA3 shows labelling of ovoid synapses (arrows) consistent with glutamate excitatory function. Myelinated axon at A. In **B**, mouse cortex again shows highly selective labeling of synapses.

**FIGURE 5:**
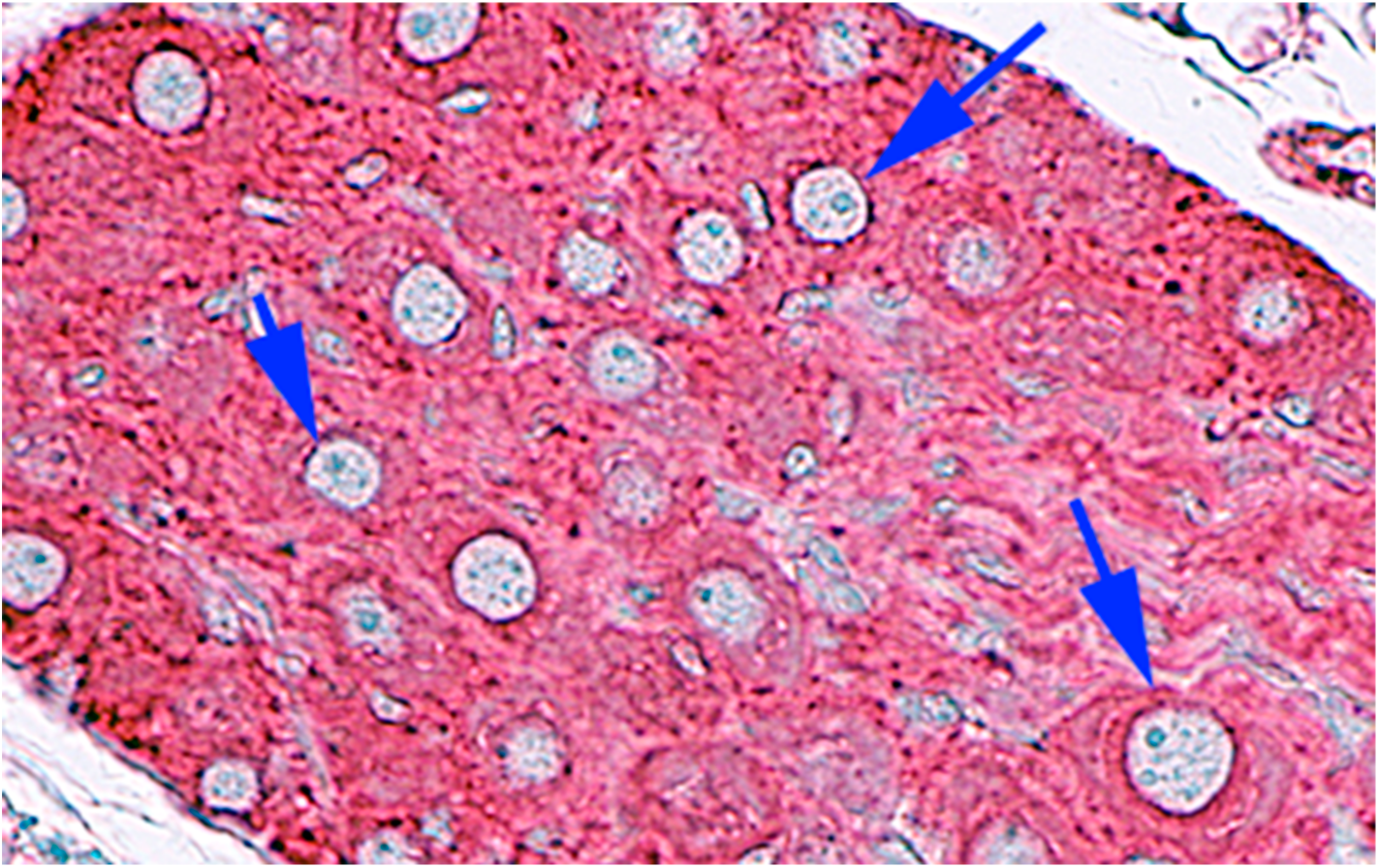
Unlike Cerebellar Purkinje neurons and Pyramidal neurons, large sympathetic ganglia neurons in the peritoneum show strongly labeled spx+ perikarya with strong perinuclear label (arrows). Tubular labeled structures in the cytoplasm are probably on the rough endoplasmic reticulum.

### Novel neurodegenerative spx pathology

In infectious TSEs, and in many other dementing neurodegenerative diseases of unknown cause, the cerebellum is often ignored as a focus of pathology. By Western blot, regions of thalamus and cortex showing CJD and scrapie terminal spongiform changes and PrP pathology failed to show any obvious quantitative or qualitative spx changes as compared to controls (25). On the other hand, normal looking brain with exponentially replicating CJD and scrapie infectious agents, but no detectable PrP pathology, have been shown to express significant molecular markers of innate immunity during the initial phases of infection. Large increases in astrocytic mRNA and myeloid/microglial activation also precede behavioral and PrP pathology (1, 6, 7). To see if we had overlooked potential changes in spx that might be regional and/or neuron-type specific, we evaluated the cerebellum of mice, hamsters, rats and Guinea pigs with proven TSE infections from a wide variety of distinct agent strains, e.g., sporadic CJD (sCJD), RML scrapie, 263k scrapie, Asiatic FU-CJD (12). Fig. 6 shows there are unique as well as shared pathological manifestations of spx in different agent disease models.

**Figure 6:**
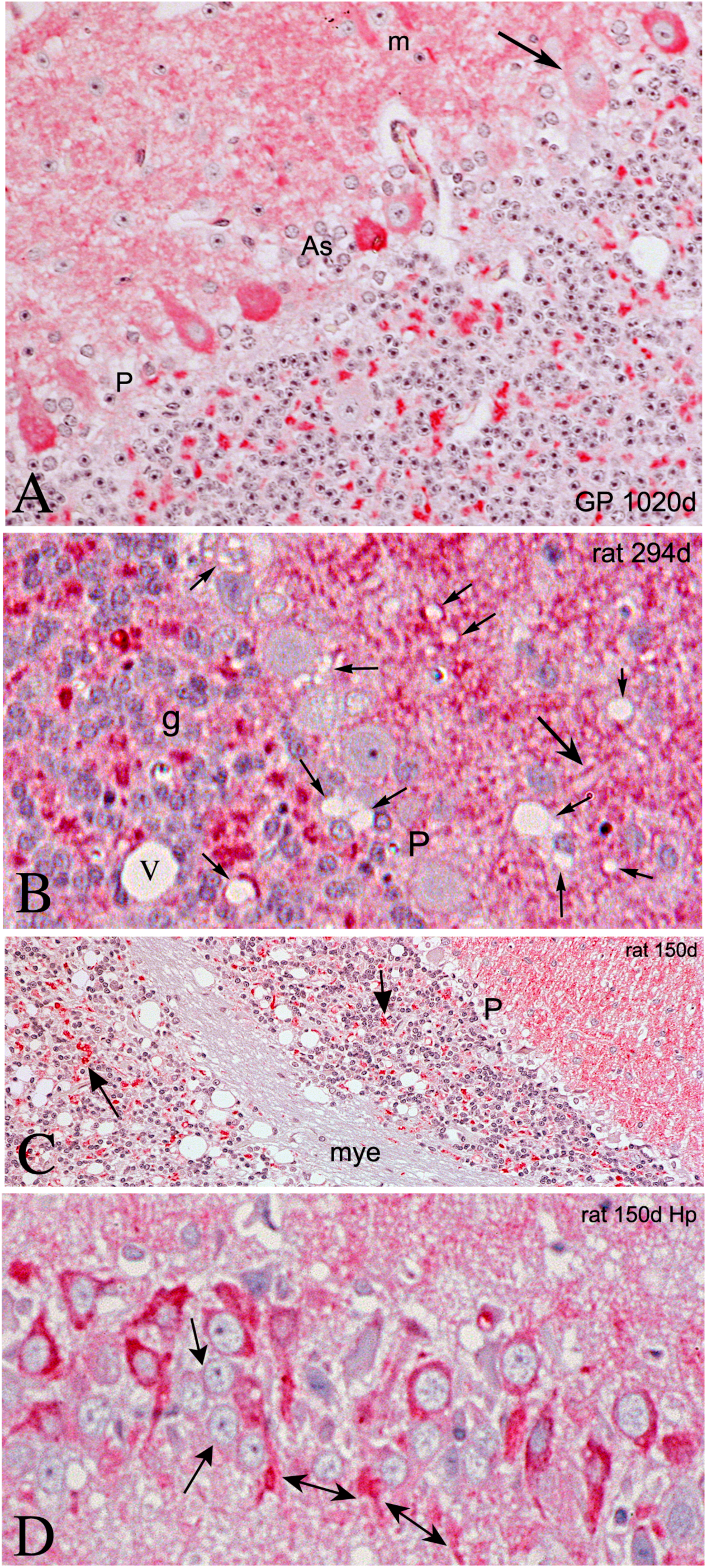
**A**: Guinea Pig with sCJD shows pathological aggregates of spx in Purkinje perikarya; some of these have a fibrillar appearance and are also seen in the dendrites around m. Only one semi-normal cerebellar Purkinje cell with low spx is seen (arrow) an others have been lost (as at P). Astrocytes are also seen where Purkinje neurons have disintegrated (As). Note the reduced number of large mossy neuron synapses in the granule cell layer and the molecular layer synapses also show less label than normal. Spongiform change (vacuoles) are rare in this ill Guinea Pig at 1020 days (<3 years old); Guinea Pigs have lived for 12 years in our lab, i.e., this is not an old animal. **B**: Clinically rat with sCJD at 294 days shows some vacuoles (small arrows) and a vessel (v) but does not show Purkinje cell spx accumulation in the perikarya. A large arrow shows normal spx+ boutons and synapses in the GC layer are numerous. **C**: in contrast to the ill rat in B, this normal appearing rat sacrificed at 150 days in the same series shows numerous vacuoles in the GC layer with dramatic loss of most mossy synapses. The myelin (mye) with axons shows negligible spx. Purkinje cells (P layer) do not show fibrillar accumulations of spx although many are missing. **D**: Instead, rat hippocampus at 150 days shows abnormal collections of spx in Pyramidal neuron perikarya extending into their dendrites. The labeled dendrites (double headed arrows) appear as fibrillary tangles and only a few more normal appearing Pyramidal neurons remain (single headed arrows).

Fig. 6A shows a Guinea Pig infected with the human sCJD agent. This clinically ill animal does not display classical TSE spongiform change in the cerebellum, yet its Purkinje cells (P) show abnormal collections of spx in the perikaryon, and these are present in all except the one remaining more normal neuron (at arrow). Spx is concentrated more at the base of the Purkinje cell as compared to the emerging apical dendrite. Moreover, these intracellular intense spx signals are reminiscent of AD neurofibrillary tangles, especially as they extend in more distal Purkinje dendrites in the molecular layer, as shown in Fig. 6A (at both sides of m). This layer also contains a few strongly positive round spx bodies representing cross-sections of degenerating dendrites. Purkinje cell neurodegeneration is ongoing, and astrocytes have collected (as at As) in regions where Purkinje cells have died and are missing. The intense bouton-like staining in the molecular layer is markedly diminished here (compare Fig. 1E). The internal granule cell layer also shows fewer large spx synaptic rosettes, indicating distal pontine excitatory neurons are degenerating. In sum, there is remarkable amount of unsuspected spx pathology in specific cells of the cerebellum that and novel, and not evident by Western blotting of heterogeneous cellular lysates.

Rats have a very different immune system than Guinea Pigs (a non-rodent species) and the sCJD agent wreaks morphologic havoc in different regions of rat brain. In Fig. 6B, an sCJD infected rat with progressive clinical signs over 40 days still maintained normal appearing Purkinje perikarya with low spx signal, unlike the Guinea Pig. This difference represents a species specific response to a single TSE agent, one that can also profoundly alter incubation time (12). This terminally ill rat at 294 days post-inoculation also displays some vacuolar degeneration in cerebellar dendrites (small arrows), but unlike the Guinea pig, multiple small synaptic boutons are still strongly positive and abundant in the molecular layer where they surround neuronal dendrites in a typical normal arrangement (as at large arrow). The large spx synapses in the GC layer are also preserved. We therefore assumed that sCJD in rats provoked little cerebellar pathology. However, in evaluating inoculated rats sacrificed well before clinical signs in this time course series (6) we were surprised to find major early pathological changes in the rat cerebellum. Fig. 6C shows a rat with no clinical signs killed at 150 days post-inoculation, i.e., ∼150 days before terminal disease of cagemates (6). There are profound vacuolar changes within the GC layer. Only very few positive spx synaptic rosettes (arrows) are seen, and this indicates early selective vacuolar disintegration of pontine mossy fibers and their synapses. Additionally, the molecular layer is only weakly positive for spx at this early time point. Notably, myelinated tracts (mye) in the cerebellum that carry the mossy fiber axons (as virtually all myelinated tracts in normal brain) also show no obvious spx signal, indicating spx may be translated and possibly transcribed in the synaptic rosette.

The early vacuolization in this cerebellum mimics the previous demonstration of TSE vacuoles elsewhere in the rat cerebrum in healthy appearing rats at 150 days, a time well before abnormal PrP responses are detected (6). It also demonstrates that spx can be useful as an early marker of neurodegeneration. This concept is furthered by the remarkable concentration and abnormal fibrillar aggregates of spx in hippocampal Pyramidal perikarya and their dendrites (double headed arrows) as early as 150 days post-inoculation (Fig. 6D). Only a few of these Pyramidal neurons may be normal (arrows). The massive abnormal spx fibrillary-like degenerative changes are not unique to Guinea Pig Purkinje cells but are also a dramatic feature of hippocampal Pyramidal neurons in a region known to have predominantly excitatory glutamate synapses. This conservation of pathology can also be useful in evaluating other diseases such as Parkinson’s Disease.

## DISCUSSION

Study of spx in the cerebellum here simplified the identification of well defined neuronal subsets with known functional attributes. The cerebellum participates in coordinating motor learning and balance, as well as other important behaviors such as cognition, emotion, and spatial navigation that are often damaged in neurodegenerative diseases. Spx showed a conserved architectural pattern in mammals. The specificity spx in synapses, with particularly high concentrations in excitatory glutamate synapses was striking and maintained in evolution, as demonstrated in the representative rodent, Guinea Pig and human cerebellar samples shown here. Previous studies have produced good evidence that this spx localization reflects a microbial phage REP segment. Binding of this spx antibody to Western blot lysates, and to various mammalian tissue cell types in major organs, was completely inhibited by inclusion of its cognate peptide (2, 25);. Both rabbit sera also decorated the same 41kDa band of REP, is in good accord with its predicted 38kDa size, and a transiently transfected REP construct studied elsewhere also appeared to be 41kDa (2, 25). Neither the REP antigenic peptide nor its corresponding complete REP DNA has been detected in any mammalian database up to the present time, and even chaotropic 2MGdnHCl treatment for enhanced penetration of antibody did not alter the fundamental pattern or conservation of spx labelling ultrastructurally. One of the most interesting facets of spx was its preferential localization in synapses as compared to much weaker signals in the perikaryon of cerebellar neurons. This was evident in GC cell bodies versus their synaptic boutons on Purkinje cell dendrites. The consistent lack of spx signal in white matter tracts that carry mossy fiber axons from the pons over long distances versus their terminal synapses on GC neurons in the cerebellum also suggests that spx is translated at the synaptic terminus. Indeed, cytoplasmic SPHINX1.8 DNA may also reside, replicate and actively transcribe in synaptic regions, like mitochondria.

Is there precedence for this unusual concept in mammalian cells? It is often implied that mitochondrial DNA replication, essential for its survival in the cytoplasm, is completely dependent on nuclear DNA functions. Nuclear encoded DNA polymerase γ, mitochondrial DNA helicase and mitochondrial single-stranded DNA-binding protein together facilitate its ss DNA replication (40). However, mitochondria have been shown to replicate in severed chick axons (37), and increases in replication in distal axons has been shown in a Parkinson Disease stress model (38). Thus SPHINX DNAs may also replicate, be transcribed, and translated in distal mammalian axons. Interestingly, the tight packing of mitochondria at mossy fiber synapses on cerebellar GC may collaborate in the processing and functions of SPHINX DNA at this site. It will be of great interest to find if other spx REPs, such as the one encoded by SPHINX 2.36 DNA also concentrate in synapses where they may enhance each others replication and functions as originally suggested (23). In any case, the persistence of the SPHINX1.8 REP at specific synapses implicates both evolutionary divergence and positive selection. This REP sequence is only is only 70% homologous to its best phage counterpart in the environment, implicating a relatively high mutation rate, as if found for mitochondria. At the same time, the REP antigenic peptide is conserved in diverse mammals, a feature that underscores its selective maintenance.

The most, fundamental and fascinating question is how and where any phage REP sequence may have entered mammals to eventually insert into the ovum. One source is the gut. Abundant gram negative bacteria, including Acinetobacter, carry as many as 10^31^ phage viruses, the most numerous biomass of the gut (41). Most of these virome sequences have not been sequenced and/or do not have sequence matches in the current database. Phages possess a unique capability of bypassing anatomical and physiological barriers, quickly permeates across the endothelium barrier, and can appear in blood [reviewed in (42)], a classic viral route of dissemination. Phages also readily penetrate the brain, especially via a nasal route. The sturdy filamentous portion of the phage, or segments of “circular nanoparticles” show even greater and prolonged penetration (42). We presume that some of these segments, such those containing special REPs, confer a selective advantage for their favor in evolution and REPs might possibly participate in cytoplasmic DNA repair. Clearly not all of the many circular DNAs in the SPHINX family are maternally inherited, and a number of other family members from cow sera and dairy products are proposed risk factors for breast and colon cancer (43). On the other hand, given the findings here, along with at least two additional SPHINX elements in brain, encourages further investigations of their role(s) in neurodegeneration and neoplasia. Complementary elements may work together combinatorically to enhance specific functions and pathology (23).

In terms of pathology, the discovery of specific spx fibril like accumulations in particular large neurons undergoing neurodegeneration are remarkable. Such changes were not entirely surprising because previous experiments here showed that ∼10% of spx protein was resistant to limited proteinase K digestion (25), a feature of abnormal PrP and other fibrillar accumulations seen in AD, and Parkinson’s disease. In TSEs, pathology in specific neuronal cell types, such as cerebellar Purkinje neurons of Guinea Pig versus hippocampal Pyramidal neurons in rat was species dependent because both were infected with the same sCJD agent. Nevertheless, the spx fibrillary change was indistinguishable and conserved within both types of neurons. Moreover, Pyramidal spx accumulations in the perikaryon and dendrites occurred ∼150 days before the clinical signs in rat sCJD suggests spx may have diagnostic use in detecting early disease. The simultaneous loss of mossy fiber synapses, with profound vacuolization in the granule cell layer of the cerebellum, also shows spx might be an early diagnostic tool; such vacuoles are concentrated ultrastructurally in dendrites and synaptic regions (44). Spx pathology further discriminated different models of CJD, and it may also define new subsets of neurodegenerative disease caused by traumatic, vascular, environmental and microbial agents.

In conclusion, the fastidious and conserved residence of spx in specific synapses, its consistent ability to highlight previously inapparent TSE agent induced changes, and its appearance early in disease, all show spx can participate in neurodegenerative processes. Future studies can use spx to target new and unsuspected pathways and targets for progressive molecular changes. On a broader evolutionary scale, these studies emphasize the importance of deeper analyses of unknown phage and microbial elements that can reside covertly in mammalian cells. Some of these sequences may initiate or cause progressive late-onset disease, especially in concert with other environmental elements or stresses (1).

## MATERIALS & METHODS

### Light microscopy preparations

Archival human and TSE experimental animal tissue were sectioned and dewaxed as previously for detection of spx using a previously characterized affinity purified rabbit antiserum (2, 25). Briefly, sections were blocked in 10% normal goat serum and 3% fish gelatin in TBS at 22°C for 1hr and then exposed to Rabbit anti-spx at 1:5,000 in TBS-0.1% Tween 20 (TBS-T) at 4°C overnight. TBS-T washes were followed by Goat anti-rabbit biotin in TBS-T at 1:5000 for 1.5 hr at 37°C, TBS-T washes and then exposed for 30min to KPL strepavidin-alkaline phosphatase and developed with Vector Red per kit instructions. For enhanced penetration of very old formalin fixed brain, heat-citrate autoclaving as described (6) or 2M GdnHCl exposure for 30 min was done after dewaxing and these treatments did not affect spx localization or peptide blocking of the anti-spx antibody as previously shown (2, 25). A Biorad confocal microscope was used to collect fluorescent optical slices which were then deconvolved using the Biorad software, and some stacks were then rotated using commercial Zen software with the help of Dr. Larry Rizzolo.

### Electron microscopy

followed a similar sequence of antibody exposure except 4% paraformaldehyde-0.1%glutaraldehye perfused mouse brains were sliced (∼50μ thick) with a vibratome and slices blocked in 3% normal goat serum, 1% BSA, 0.1% lysine, 0.1% glycine for 24 hr prior to exposure for 2 days to anti-spx antibody followed by biotinylated anti-rabbit antibody (Vector 1:250) and Avidin-Biotin peroxidase (Vector 1:200) that was visualized with Diaminobenzidine. Sections were exposed to 1% osmium tetroxide and dehydrated in increasing ethanol concentrations. I% uranyl acetate was added to the 70% ethanol to enhance ultrastructural membrane contrast. Flat embedding in Durcupan followed dehydration. Ultrathin sections were cut on Leica ultramicrotome, collected on formvar-coated single slot grids, and analyzed with Tecnai 12 Biotwin Electron Microscope. For enhanced penetration, slices were equilibrated in 20% glycerol and freeze-thawed x3 or exposed to 2MGdnHCl for 8-10 min before blocking. Both treatments increased the number of synapses labeled but did not alter the pattern of labeling as shown representatively in Fig. 3A,B; these were the same as with TBS-T. Sections were osmified, embedded in plastic and thin sectioned for electron microscopy.

## References

1. Manuelidis L (2013) Infectious particles, stress, and induced prion amyloids: a unifying perspective. Virulence 4(5):373–383.

2. Manuelidis L (2019) Prokaryotic SPHINX 1.8 REP protein is tissue-specific and expressed in human germline cells. J Cell Biochem 120(4):6198–6208.

3. Manuelidis L (1994) Dementias, neurodegeneration, and viral mechanisms of disease from the perspective of human transmissible encephalopathies. Ann NY Acad. Sci 724:259–281.

4. Kipkorir T, Tittman S, Botsios S, & Manuelidis L (2014) Highly Infectious CJD Particles Lack Prion Protein but Contain Many Viral-Linked Peptides by LC-MS/MS. J Cell Biochem 115(11):2012–2021.

5. Botsios S & Manuelidis L (2016) CJD and Scrapie Require Agent-Associated Nucleic Acids for Infection. J Cell Biochem 117(8):1947–1958.

6. Manuelidis L, Fritch W, & Xi YG (1997) Evolution of a strain of CJD that induces BSE-like plaques. Science 277:94–98.

7. Lu ZH, Baker C, & Manuelidis L (2004) New molecular markers of early and progressive CJD brain infection. J. Cellular Biochem. 93:644–652.

8. Shlomchik MJ, Radebold K, Duclos N, & Manuelidis L (2001) Neuroinvasion by a Creutzfeldt-Jakob disease agent in the absence of B cells and follicular dendritic cells. Proc Natl Acad Sci U S A 98(16):9289–9294.

9. Prusiner SB (1982) Novel proteinaceous infectious particles cause scrapie. Science 216:136–144.

10. Prusiner S (1998) The Nobel Lecture: Prions. Proc Natl Acad Sci U S A. 95(23):13363–13383.

11. Telling G, et al. (1996) Evidence for the conformation of the pathologic isoform of the prion protein enciphering and propagating prion diversity. Science 274:2079–2082.

12. Manuelidis L, Chakrabarty T, Miyazawa K, Nduom NA, & Emmerling K (2009) The kuru infectious agent is a unique geographic isolate distinct from Creutzfeldt-Jakob disease and scrapie agents. Proc Natl Acad Sci U S A 106(32):13529–13534.

13. Arjona A, Simarro L, Islinger F, Nishida N, & Manuelidis L (2004) Two Creutzfeldt-Jakob disease agents reproduce prion protein-independent identities in cell cultures. Proc Natl Acad Sci USA 101:8768–8773.

14. Manuelidis L & Lu ZY (2003) Virus-like interference in the latency and prevention of Creutzfeldt-Jakob disease. Proc. Natl. Acad. Sci. USA 100: 5360–5365.

15. Nishida N, Katamine S, & Manuelidis L (2005) Reciprocal interference between specific CJD and scrapie agents in neural cell cultures. Science 310:493–496.

16. Marin-Moreno A, Aguilar-Calvo P, Pitarch JL, Espinosa JC, & Torres JM (2018) Nonpathogenic Heterologous Prions Can Interfere with Prion Infection in a Strain-Dependent Manner. J Virol 92(24).

17. Legname G, et al. (2004) Synthetic mammalian prions. Science 305:673–676.

18. Wang F, Wang X, Yuan C-G, & Ma J (2010) Generating a prion with bacterially expressed recombinant prion protein. Science 327:1095–9203.

19. Timmes AG, Moore RA, Fischer ER, & Priola SA (2013) Recombinant prion protein refolded with lipid and RNA has the biochemical hallmarks of a prion but lacks in vivo infectivity. PLoS One 8(7):e71081.

20. Schmidt C, et al. (2015) A systematic investigation of production of synthetic prions from recombinant prion protein. Open Biol 5(12):150165.

21. Barron RM, et al. (2016) PrP aggregation can be seeded by pre-formed recombinant PrP amyloid fibrils without the replication of infectious prions. Acta neuropathologica 132(4):611–624.

22. Manuelidis L, Yu Z-X, Barquero N, & Mullins B (2007) Cells infected with scrapie and Creutzfeldt-Jakob disease agents produce intracellular 25-nm virus-like particles. Proc Natl Acad Sci USA 104:1965– 1970.

23. Manuelidis L (2011) Nuclease resistant circular DNAs copurify with infectivity in scrapie and CJD. J Neurovirol 17(2):131–145.

24. Whitley C, et al. (2014) Novel replication-competent circular DNA molecules from healthy cattle serum and milk and multiple sclerosis-affected human brain tissue. Genome announcements 2(4).

25. Yeh YH, Gunasekharan V, & Manuelidis L (2017) A prokaryotic viral sequence is expressed and conserved in mammalian brain. Proc Natl Acad Sci U S A 114(27):7118–7123.

26. Cajal SR (1955) *Histologie du Systeme Nerveux de L’homme & des vetebres* (Instituto Ramon y Cajal, Madrid Spain); trans Azoulay L.

27. Masoli S & D’Angelo E (2017) Synaptic Activation of a Detailed Purkinje Cell Model Predicts Voltage-Dependent Control of Burst-Pause Responses in Active Dendrites. Frontiers in cellular neuroscience 11:278.

28. Einstein G, Buranosky R, & Crain BJ (1994) Dendritic pathology of granule cells in Alzheimer’s disease is unrelated to neuritic plaques. J Neurosci 14(8):5077–5088.

29. Sagan L (1967) On the origin of mitosing cells. J Theor Biol 14(3):255–274.

30. Rosario K, Schenck RO, Harbeitner RC, Lawler SN, & Breitbart M (2015) Novel circular single-stranded DNA viruses identified in marine invertebrates reveal high sequence diversity and consistent predicted intrinsic disorder patterns within putative structural proteins. Frontiers in microbiology 6:696.

31. Manuelidis L (1982) Nucleotide sequence definition of a major human DNA, the Hind III, 1.9 kb family. Nucleic Acids Res. 10:3211–3219.

32. Manuelidis L (1990) A view of interphase chromosomes. Science 250:1533–1540.

33. Taruscio D & Manuelidis L (1991) Integration site preferences of endogenous retroviruses. Chromosoma 101:141–156.

34. Conley AB, Piriyapongsa J, & Jordan IK (2008) Retroviral promoters in the human genome. Bioinformatics 24(14):1563–1567.

35. Grandi N & Tramontano E (2018) Human Endogenous Retroviruses Are Ancient Acquired Elements Still Shaping Innate Immune Responses. Front Immunol 9:2039.

36. Sotelo C & Llinas R (1972) Specialized membrane junctions between neurons in the vertebrate cerebellar cortex. J Cell Biol 53(2):271–289.

37. Amiri M & Hollenbeck PJ (2008) Mitochondrial biogenesis in the axons of vertebrate peripheral neurons. Dev Neurobiol 68(11):1348–1361.

38. Van Laar VS, et al. (2018) Evidence for Compartmentalized Axonal Mitochondrial Biogenesis: Mitochondrial DNA Replication Increases in Distal Axons As an Early Response to Parkinson’s Disease-Relevant Stress. J Neurosci 38(34):7505–7515.

39. Le Duigou C, Simonnet J, Telenczuk MT, Fricker D, & Miles R (2014) Recurrent synapses and circuits in the CA3 region of the hippocampus: an associative network. Frontiers in cellular neuroscience 7:262.

40. Ciesielski GL, Oliveira MT, & Kaguni LS (2016) Animal Mitochondrial DNA Replication. Enzymes 39:255–292.

41. De Sordi L, Lourenco M, & Debarbieux L (2019) “I will survive”: A tale of bacteriophage-bacteria coevolution in the gut. Gut Microbes 10(1):92–99.

42. Huh H, Wong S, St Jean J, & Slavcev R (2019) Bacteriophage interactions with mammalian tissue: Therapeutic applications. Adv Drug Deliv Rev.

43. H ZH, Bund T, & EM dV (2019) Specific nutritional infections early in life as risk factors for human colon and breast cancers several decades later. Int. J. Cancer 144:1574–1583.

44. Manuelidis E, Kim J, Angelo J, & Manuelidis L (1976) Serial propagation of Creutzfeldt-Jakob disease in guinea pigs. Proc. Natl. Acad. Sci. (USA) 73:223–227.

